# Sleep Enhances Neural Representations of True and False Memories: An Event-Related Potential Study

**DOI:** 10.1101/2020.10.21.349530

**Authors:** Sophie Jano, Julia Romeo, Matthew D. Hendrickx, Alex Chatburn

## Abstract

Episodic memory is reconstructive and is thus prone to false memory formation. Although false memories are proposed to develop via associative processes, the nature of their neural representations, and the effect of sleep on false memory processing is currently under debate. The present research employed the Deese-Roediger-McDermott (DRM) paradigm and a daytime nap to determine whether semantic false memories and true memories could be differentiated using event-related potentials (ERPs). We also sought to illuminate the role of sleep in memory formation and learning, with the use of a daytime nap. Healthy participants (*N* = 34, 28F, mean age = 23.23, range = 18-33) completed the DRM task with the learning and recognition phases separated by either a 2hr daytime nap or an equivalent wake period. Linear mixed modelling revealed larger LPC amplitudes for true memories in contrast to false memories, and larger P300 amplitudes for false compared to true memories. Those in the nap group also exhibited larger LPC and P300 amplitudes than participants in the wake group. Additionally, a larger P300 amplitude at delayed recognition (following a consolidation opportunity) was associated with increased true memory accuracy. These findings are argued to be reflective of sleep’s ability to promote pattern separation and pattern completion, with true memories arising from distinct memory traces, and false memories arising from thematic extraction and overlap in neural representations. The present research supports the perspective that both true and false memories are reflective of adaptive memory processes, whilst also suggesting that P300 amplitude affects episodic memory accuracy.

## Introduction

Memory is often misconceived as a concrete record of experience, prompting the belief that it is accurate and infallible (Diekelmann et al., 2008). Contrary to popular opinion, research has demonstrated that memory is a reconstructive process that is highly susceptible to distortion, inaccuracies, and generalisation across both wake and sleep (Diekelmann et al., 2008). This flexibility renders memory important for learning, with generalisation and distortion providing insight into the brain’s associative processes.

The reconstructive nature of memory is strongly demonstrated by false memory paradigms, which leverage semantic associative processing to induce memories for events or details of events that have not occurred (Straube, 2012). False memories can arise at multiple stages of the memory process (encoding, sleep-based consolidation, and retrieval), with their inaccuracy passing below the owner’s conscious awareness, thus rendering them difficult to detect (Abe et al., 2008).

Although false memories can be reflective of memory errors, they can also be indicative of memory association, depending on the task of focus. Therefore, they reflect an adaptive underlying process, although the complete nature of this effect in the brain is unclear (Deese, 1959). Distinguishing between true and false memories using neurophysiological techniques could provide insight into whether they reflect the same or differing underlying processes, potentially enabling the discrimination of learning and memory in the brain. The purpose of the current research was to investigate whether true (veridical) and false (generalised) memories can be differentiated using neuroscientific techniques, and if so, to determine the role of sleep in false memory generation. To address this question, an investigation of semantic false memory and a daytime nap was conducted, with the use of the Deese-Roediger McDermott (DRM) paradigm and the event-related potential (ERP) method.

### Semantic False Memory

A semantic false memory occurs when an individual recalls stimuli or events that share semantic commonalities with the original learned inputs, but which were never originally encountered (Diekelmann et al., 2009). This is commonly observed in the DRM paradigm, wherein participants falsely recognise a related critical lure word (e.g. *sleep*) as comprising an earlier presented list of related study words (e.g. *bed, dream, wake, blanket etc.*), despite its absence from that original list (Deese, 1959; Jou et al., 2017; Roediger & McDermott, 1995). Such memories involve the false recognition of related content, and are posited to rely on associative networks, whereby the essential elements (the “gist”) of the memory trace are integrated with existing, related memories (Straube, 2012). This may be complementary to predictive coding accounts, which propose that the brain utilises top-down processing to construct generative schemata of the world, thus enabling it to make predictions about incoming sensory information (Aitchison & Lengyel, 2017; Heilbron & Chait, 2017). According to predictive coding, semantic false memories may arise due to the generalisation of specific, related content into a common, broader schema, which is subsequently applied during recognition to minimise processing effort (Aitchison & Lengyel, 2017).

Although semantic false memories may arise from predictive processes, these may manifest in multiple ways. In line with a recent two-process theory, they may be either expected (because the lure words that elicit them fit well with the broader theme; thus leading to diminished behavioural and/or neural responses to stimuli), or surprising (because such words are not part of the original list, leading to increased neurobehavioral responses; Press et al., 2020). This uncertainty could be resolved with the use of neuroscientific techniques such as electroencephalography (EEG), which could also provide insight into how expectancy and surprise may influence experience-based decision making.

### Event-Related Potentials (ERPs)

Event-related potentials (ERPs) have rarely been used to determine whether true and false semantic memories are distinguishable at a neural level, potentially because false memory paradigms typically lack the large number of trials necessary for the calculation of ERPs (Beato & Díez, 2011). Of the few studies that have used ERPs to investigate DRM effects, some have observed larger amplitudes in the late positive component (LPC; a parietal ERP arising 400 to 800ms post-stimulus onset) for true memories in contrast to false memories (Chen et al., 2012; Curran, 2000; Curran et al., 2001; Nessler et al., 2001). This is proposed to reflect a greater engagement of recognition processes, which involve the retrieval of detailed episodic information (Yonelinas, 2002).

A late frontal positivity (900ms to 1500ms post stimulus onset) has also exhibited significantly larger amplitudes for true compared to false memories (Curran, 2000; Geng et al., 2007; Nessler et al., 2001; Wiese & Daum, 2006). However, as previous research has observed lower test-retest reliabilities for late, as opposed to early components (Kompatsiari et al., 2016), the reliability of the late frontal component may be questionable.

Findings regarding the LPC suggest that semantic false memories may differ from true memories at an electrophysiological level (Curran, 2000; Wiese & Daum, 2006). However, this is contradicted by other DRM studies which have failed to locate ERP differences between the two, raising the alternative suggestion that both may share common mechanisms (Beato et al., 2012; Boldini et al., 2013; Curran et al., 2001). This may be harmonious with predictive coding, because if all related content is generalised into broader schemata, it is possible that true and false memories would be represented similarly in the brain (Aitchison & Lengyel, 2017).

Another ERP that could shed light upon the neural underpinnings of true and false memories is the P300. The P300 ERP is typically maximal over parietal sites and is elicited during ‘Oddball’ paradigms, in response to infrequent target stimuli, interweaved amongst frequent, irrelevant stimuli (Campanella et al., 2010; Linden, 2005). This has prompted the widespread understanding of the P300 as a neural index of target identification and goal-directed attention (Linden, 2005). As a DRM-based semantic false memory arises from a lure in which the participant is actively searching for previously seen words (targets), it is suggested that a P300 effect may be present (Linden, 2005). However, in the context of false memory, the pattern in which this component may appear is unclear, as the literature on the P300 for discriminating between true and false memories is limited and has achieved mixed results (Allen & Mertens, 2009; Miller et al., 2001). This highlights the need for further research on the neural distinctions between the two types of memory.

### A Potential Role for Sleep

The effect of sleep on semantic false memory is also under-researched using ERPs, despite decades of accumulated evidence for the influence of sleep on memory consolidation (Diekelmann, et al., 2009; Rasch & Born, 2008). Previous DRM and sleep literature has employed naps, which are effective in investigating sleep functions as they have similar characteristics to a full night of sleep (Durrant et al., 2011; Shaw & Monaghan, 2017). The few studies conducted to date have associated daytime naps with an increased number of DRM false memories, compared to wake conditions (Payne et al., 2009; Shaw & Monaghan, 2017). This, in accordance with predictive coding, may suggest that sleep facilitates the integration of information into broader schemata (Heilbron & Chait, 2017; Lewis & Durrant, 2011). The contextual binding (CB) approach (Yonelinas et al., 2019) may also extend upon this idea, as it posits that slow wave sleep (SWS) facilitates the binding of item information with contextual information, and thus improves memory. By considering the broader theme of a DRM list as the context of the DRM words, it could be suggested that sleep enhances the binding between the related theme and the word of focus (Stadler et al., 1999; Yonelinas et al., 2019). If the word has more strongly been bound to its broader context, other contextually related words (e.g. related critical lures) are more likely to seem familiar, possibly resulting in a false memory (Yonelinas et al., 2019).

Therefore, contexts in CB can be interpreted similarly to the predictive coding idea of a model, with the CB approach providing a more specific mechanistic interpretation of how predictive coding may arise in the brain (Yonelinas et al., 2019). Additionally, if the models/contexts are more generalised in nature, it could be suggested that sleep would lead to lower LPC amplitudes for false memories, which are understood to be more generalised, as opposed to true memories, which are understood to be more episodic (Straube, 2012). However, this is yet to be investigated directly. This suggestion is also contradicted by other research demonstrating higher false memory rates in sleep-deprived participants (Chatburn et al., 2016; Diekelmann et al., 2008; Fenn et al., 2009). Therefore, a deeper investigation of sleep’s role in false memory generation, and the neural underpinnings of this relationship, is needed.

### Research Aims and Hypotheses

This study aimed to illuminate the underlying processes of semantic false memory, firstly by determining whether ERPs can differentiate between true and false memories. In terms of learning, such an investigation provides insight into how knowledge can be created from discrete units of data and of how the brain generalises across experience to extract the broader gist and predict incoming sensory information.

The current research combined ERPs and the DRM paradigm, with the DRM immediate and delayed recognition phases being separated by either a nap or wake period. This enabled a direct comparison of the ERPs of false and true memories, and of the effects of sleep compared to wake. Based on the previous ERP literature, it was predicted that:

Hypothesis 1: *True memories would be associated with larger LPC amplitudes than semantic false memories.*

To determine how sleep affects the neural representations of true and false memories, the following research question was proposed:

Research question 1: *Does sleep affect LPC amplitude for true and false memories?*

Due to the evidence for sleep’s role in the generalisation of information, it was also hypothesised that:

Hypothesis 2*: Participants in the nap group will exhibit greater behavioural endorsements of semantic false memories, than participants in the wake group.*

Finally, as the P300 may have implications for understanding the relationship between goal-directed attention and false memories, we also asked:

Research Question 2: *Does P300 amplitude differ between true and false memories?*

## Method

### Participants and Recruitment

52 people expressed interest in the study, with 39 meeting the eligibility criteria. All individuals that took part in the research were healthy, right-handed adults between the ages of 18 and 45, who were not taking any regular medication that could interfere with the EEG, had not taken recreational drugs in the previous 6 months, and who did not have a diagnosed psychiatric or sleep disorder. Healthy participants were chosen to avoid confounding effects from other conditions, and the age group was selected to prevent the EEG from being affected by neural differences pertaining to age-related cognitive decline. Three participants were excluded due to technical issues and another was excluded due to signal noise in the EEG. As a result, a total of 34 participants (28 females, 6 males, mean age = 23.23, standard deviation = 4.19) were included in the final data analysis, with 18 in the experimental (daytime nap) group, and 16 in the control (wake) group.

Ethical approval for the experiment was acquired from the UniSA Human Research Ethics Committee, and all participants provided written informed consent. Participants also received a $40 honorarium for their time. A G*Power calculation based on previous DRM sleep and memory studies by Diekelmann et al. (2008) revealed that in order to obtain sufficient power (0.80) to detect a large effect size (0.80) at a significance level of .05, 30 participants were recommended (Faul et al., 2007).

### Measures

#### The Pittsburgh Sleep Quality index (PSQI)

The PSQI is a self-report questionnaire designed to measure sleep quality over the past month (Buysse et al., 1989). It is typically used as a screening tool for poor sleep in non-clinical adult populations (Mollayeva et al., 2016). It was employed in the present research to screen for and exclude those with poor sleep quality, as this may have influenced the participants’ ability to nap or their general cognitive performance (Gobin et al., 2015).

The PSQI consists of 19 items that measure seven sleep components; subjective sleep quality, sleep latency, sleep duration, habitual sleep efficiency, sleep disturbances, use of sleeping medications, and daytime dysfunction (Bush et al., 2012; Buysse et al., 1989). Scores for each component range from 0 to 3 and are summed to produce a total score ranging from 0 to 21. Scores greater than 5 are considered to reflect poor sleep quality (Alsaadi et al., 2013).

Research has demonstrated that the PSQI has high internal consistency (α=.80) and high test-retest reliability (r= .87) in intervals of two days and several weeks, suggesting that it obtains consistent results across time (Backhaus et al., 2002; Carpenter & Andrykowski, 1998). The PSQI has also demonstrated high correlations with sleep log data, suggesting that it also has high convergent validity (*r*= 0.71 to 0.81) and is likely to be measuring sleep quality (Backhaus et al., 2002). The PSQI took approximately 5 minutes to complete and participants with scores above 5 were excluded from the research.

#### The Flinders Handedness survey (Flanders)

In keeping with standard EEG practice, the current research excluded left-handed individuals in an effort to eliminate potential handedness effects (Nicholls et al., 2013). Participants completed the Flanders, which is a self-report measure of hand preference, in order to confirm right-handedness. The Flanders consists of 10 items to which participants respond “left,” “either” or “right.” Each response is weighted −1, 0 and +1 respectively, and responses are summed to obtain an overall score. Scores ranging from −10 to −5 indicate predominant left-handedness, scores from −4 to 4 indicated mixed-handedness, and scores from 5 to 10 are indicative of right-handedness (Nicholls et al., 2013). The Flanders has very high internal consistency (α= .96), however the research lacks a measure of validity as the test is relatively new (Nicholls et al., 2013). The Flanders took approximately 2 minutes to complete.

#### Physiological measures: Electroencephalography (EEG)

Participants’ EEG was recorded with a 32-channel DC BrainAmp (Brain Products GmbH), in conjunction with a bipolar electrooculogram (EOG). EOG electrodes were placed 1cm away from the outer canthus of each eye diagonally, in accordance with typical sleep EEG protocol. The EEG voltage was sampled at a rate of 1000Hz, and the impedances of all electrodes were maintained below 10kΩ. The reference electrode was FCz and the ground electrode was AFz. EEG was used in the present study to record participants’ brain activity during the DRM task and the nap period, for the calculation of ERPs and sleep staging.

#### Semantic false memory task: The Deese-Roediger-McDermott paradigm (DRM)

The DRM paradigm is a word-learning task designed to elicit semantic false memories (Roediger & McDermott, 1995). The current research employed a modified version of the typical DRM paradigm, which was presented via the *OpenSesame* software program (Mathôt et al., 2012) and which consisted of 18 lists. Previous research has indicated that the 18 lists that were used in the current experiment reliably elicit false memories 77% of the time (Stadler et al., 1999). Split-half correlations of the DRM paradigm have also revealed high internal consistency (r= .80) (Stadler et al., 1999). However, measures of convergent validity are lacking as comparable paradigms do not exist.

A typical DRM task consists of 12-15 lists, each comprised of 15 study words and one critical lure word (Roediger & McDermott, 1995). The study words are ordered in terms of strength of association to the critical lure, ranging from most related, to least related (Roediger & McDermott, 1995). However, as ERPs are created by averaging over multiple trials for each condition, the present task employed 18 lists, each comprised of 14 related words and 2 critical lures, in order to increase the number of trials and reduce the signal to noise ratio (Beato & Díez, 2011). The first word in each original list was used as the additional critical lure; a practice that has previously been shown to be effective in eliciting semantic false memories and in improving the quality and accuracy of the ERPs (Beato & Díez, 2011).

### Procedure

The procedure for the present study is presented in Figure 1. Interested participants completed a demographic questionnaire, the Flanders and the PSQI to confirm their eligibility. If eligible, they were invited to take part in the research. Prior to the day of testing, participants were asked to restrict their usual sleep time by one hour, to increase their chances of sleeping during the nap period. This was measured through self-report.

**Figure 1.**
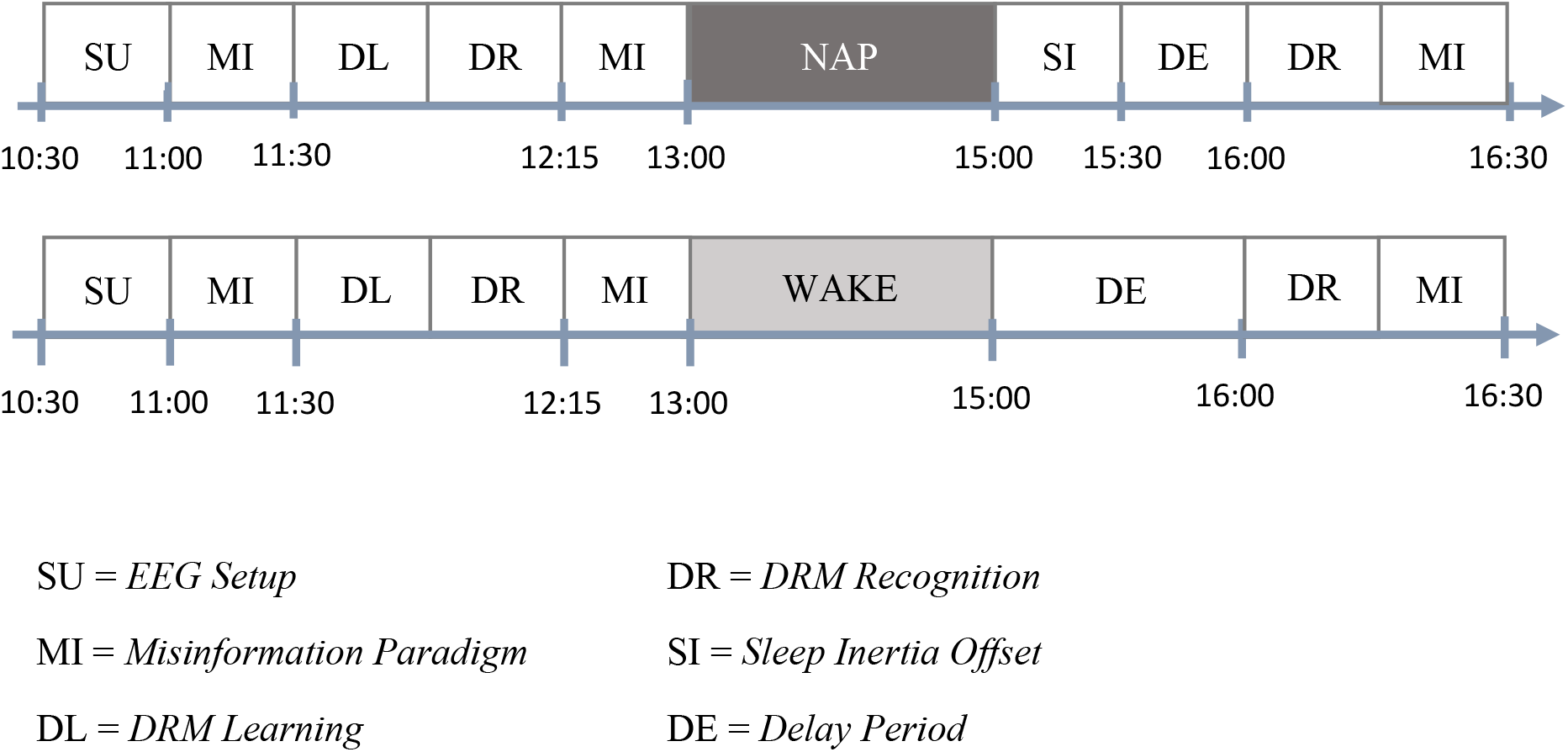
Experimental protocol of testing session. This study was part of a larger experiment that included a misinformation false memory paradigm that was unrelated to the present investigation.

#### Learning

Testing commenced at 10:30am, where participants signed a written consent form and were fitted with an EEG cap. They were then seated in front of a computer in a quiet testing room, where they completed the learning phase of the DRM task. The learning phase involved the presentation of all 14 study words from the 18 experimental lists (excluding the critical lures) in a serial visual presentation format (See Figure 2a). Each word was presented for 1250ms, and was followed by a 250ms blank interstimulus interval, before a fixation dot was presented for 500ms. Participants were instructed to learn the word lists to the best of their ability. Learning took approximately 20 minutes and participants’ EEG was recorded throughout.

**Figure 2.**
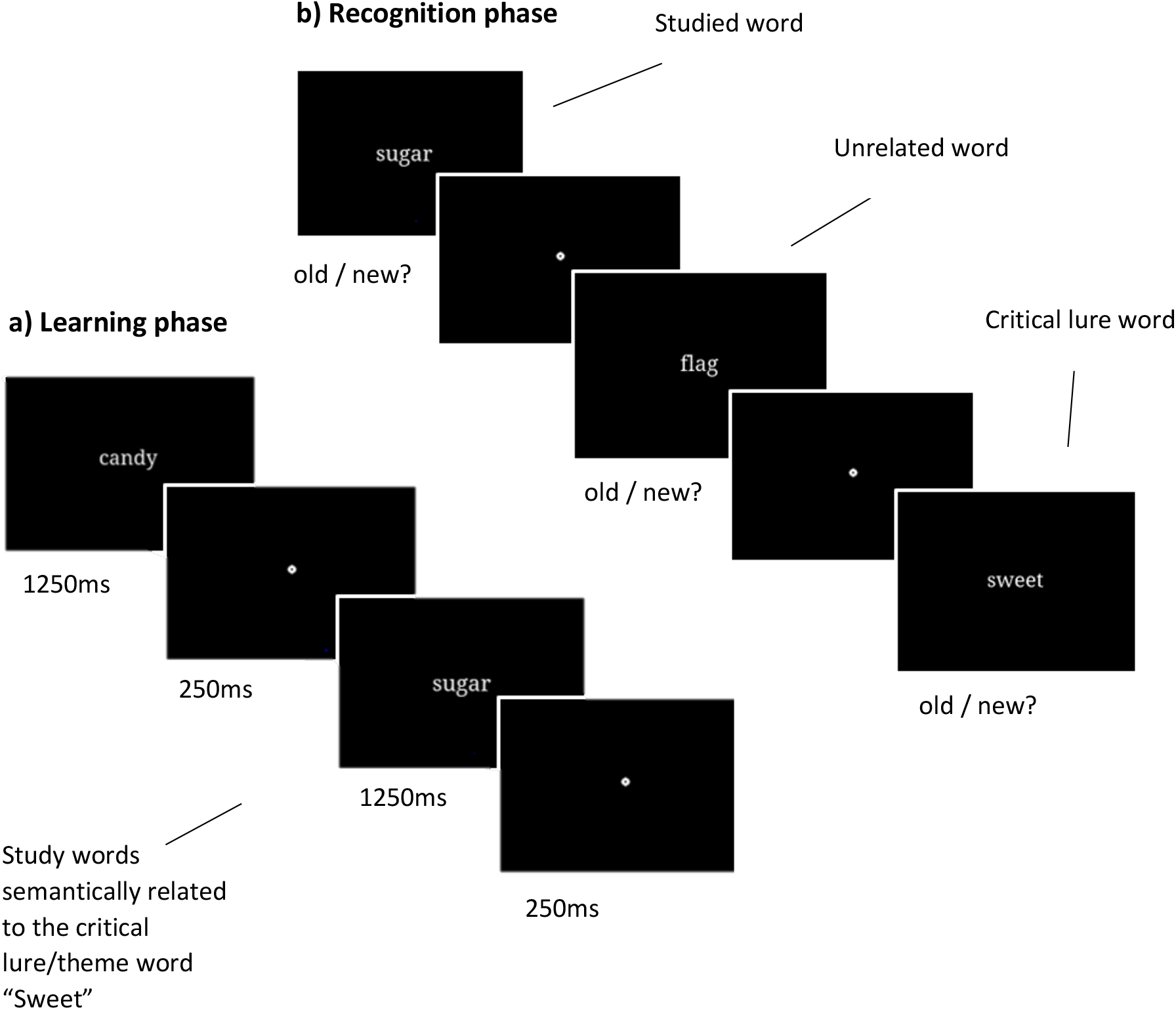
Example of the DRM paradigm. **a)** The learning phase involved viewing lists of related study words and committing them to memory. **b)** The recognition phases included immediate and delayed recognition, which involved the presentation of study words, their respective lures, and control words, to which participants made a self-paced old/new judgement.

#### Immediate recognition (IR)

Following learning, participants undertook the immediate recognition (IR) phase while their EEG was recorded. Participants viewed sub-lists consisting of the first, fifth and eighth word from each original list, plus the two critical lures, on a computer screen (See Figure 2b). The first, fifth and eighth words from unrelated control lists were also presented, in conjunction with their 2 respective lures. All words were presented in the same format and timing as in the learning phase; however, their order was pseudo-randomised to ensure that no two words from the same list would be adjacent. In response to each word, participants were instructed to make a self-paced old/new judgement using a keyboard press. “old” judgements indicated that the word was seen in the original learning phase and “new” judgements specified that it was not. Such judgements enabled the following four types of outcomes to be calculated for true memories:

- Hit: an old word is correctly identified as old
- Miss: an old word is incorrectly identified as new
- Correct rejection: a distractor/control word is correctly identified as new
- False alarm: a distractor/control word is incorrectly identified as old

It also enabled the following four outcomes for false memories:

- Hit: a critical lure word is identified as old
- Miss: a critical lure word was not identified as old
- Correct rejection: a distractor/control lure is correctly identified as new
- False alarm: a distractor/control lure is incorrectly identified as old

The separation of true and false memories, such that each was a distinct target behaviour with its own hits, misses, correct rejections and false alarms facilitated the calculation of d’, with both true and false memories receiving a separate d’ score (McNicol, 2005). It also coincided with the theoretical position that both true and false memories are not mutually exclusive, but instead represent distinct but complementary memory processes (Straube, 2012).

#### Sleep/wake

Following IR, participants were randomly allocated to either the experimental (daytime nap) group, or to the control (wake) group. Participants in the experimental group slept in the lab for a two-hour period under controlled conditions (see Figure 2). Their EEG was recorded throughout, and a nap was defined as any sleep that reached stage 2 sleep onwards, as prior sleep stages are reflective of a transitory phase between sleep and wake (Rasch & Born, 2008). A two-hour period was chosen to provide participants with enough time to fall asleep and enter at least stage 2 sleep. The nap took place at approximately 01:00pm, when circadian sleep pressure is typically high (Monk, 2005). In contrast, those in the control group engaged in tasks of low cognitive demand in the lab for two hours while maintaining wakefulness. Their EEG was not recorded during this time.

#### Delayed recognition (DR)

After the two-hour retention interval, all participants undertook a one-hour break where they engaged in tasks of low cognitive demand. The purpose of the first half of the break was to allow for the sleep inertia of those in the experimental groups to pass, whilst the second half was included to maintain consistent timing with the other groups included in the wider experiment (Achermann & Borbély, 1994). Finally, all participants completed the DRM delayed recognition phase, where they were presented with the same words as in the immediate recognition phase and were again instructed to make an old/new judgement. The presentation and timing of words was identical to that of IR and learning, although words were pseudo-randomised again to reduce order effects. Participants’ EEG was recorded throughout, and both IR and DR each took approximately 15 minutes to complete.

### Data Analysis

#### Mean ERP peak amplitude

The MNE toolbox in Python (Gramfort et al., 2013) was used to filter and pre-process the EEG data, and to remove ocular artefacts. The EEG data was filtered with a high-pass filter at 0.1Hz, and a low-pass filter at 30Hz. ERP peak amplitudes were calculated from the EEG activity arising at immediate and delayed recognition in response to the correct recognition of true memories and endorsements of false memories. Time windows that ranged from 150ms pre-stimuli onset, to 350ms post-stimuli onset, were analysed for the P300. To explore differences in the LPC, a time window from 400ms to 800ms was analysed. ERP analyses were completed over centro-medial regions for the LPC, and over posterior-medial regions for the P300, in accordance with the previous literature (Campanella et al., 2010; Chen et al., 2012)

#### DRM recognition accuracy

DRM recognition accuracy was calculated from participants’ old/new judgements to experimental words, control words, lures and control lures in the IR and DR tasks. Both true and false memories received a validated sensitivity index of memory performance (d’), adapted from signal detection theory (McNicol, 2005). d’ for both true and false memories was calculated using the program *R*, version 3.6.1 (R Core Team, 2018), with the package *Psycho* (Makowski, 2018), which calculates the Z-score of the hit-rate minus the Z-score of the false-alarm rate. Higher d’ scores for true memories indicated a greater amount of true memories, whilst higher scores for false memories indicated a greater amount of false memories.

#### Sleep

The sleep EEG data was scored by an experienced technician in 30 second epochs, in accordance with the American Academy of Sleep Medicine (AASM) guidelines (Berry et al., 2012). Information regarding total sleep time, sleep onset latency (SOL), and the percentage of time spent in sleep stages N1, N2, S3 and rapid eye movement sleep (REM) was acquired.

#### Statistical analysis

Five linear mixed-effects models (LMM) fit by maximum likelihood were calculated with the *lme4* package (Bates et al., 2014) in *R* version 3.6.1 (R Core Team, 2018). Model structures included condition, memory type, task (IR/DR), pre-stimulus LPC amplitude, pre-stimulus P300 amplitude, LPC amplitude and P300 amplitude as fixed effects. Subject ID was included as a random effect in all models, in order to account for between-participant variance (Fox, 2015). P-value estimations were constructed using Type II Wald χ2-tests from the *car* package (Fox et al., 2012), and effects were plotted using *ggplot2* (Wickham, 2016).

## Results

A summary of participants’ recognition accuracy rates for the DRM are reported in Table 1 in the form of average d’ scores. Averaging of the false memory endorsement rates per participant revealed an overall false memory endorsement rate of 72% across IR and DR, and across conditions, suggesting adequate strength of the DRM effects. All participants performed above chance, suggesting that they were engaging with the DRM task.

**Table 1.** M (SD) of d’ scores across conditions, tasks and types of memory.

| Condition | Immediate Recognition |  | Delayed Recognition |  |
| --- | --- | --- | --- | --- |
|  | True Memories | False Memories | True Memories | False Memories |
| Sleep | 1.53 (0.65) | 1.23 (0.37) | 1.24(0.65) | 1.23(0.54) |
| Wake | 1.50 (0.43) | 1.22 (0.43) | 1.15 (0.44) | 1.15 (0.56) |

### Sleep Parameters

Sleep parameters are presented in Table 2. All participants slept during the designated period, with a mean sleep duration across participants of 93.47 minutes.

**Table 2.** Means, standard errors and ranges for sleep parameters.

| Sleep parameters |  |  |
| --- | --- | --- |
|  | Mean (SEM) | Range |
| TST (mins) | 93.47 (5.9) | 28 – 115 |
| SOL (mins) | 16.08 (3.15) | 4 – 35.5 |
| N1 (%) | 18.27 (3.16) | 3.2 – 28.7 |
| N2 (%) | 54.39 (5.05) | 24.3 – 76.9 |
| SWS (%) | 9.20 (3.54) | 0 – 25.9 |
| REM (%) | 9.72 (3.54) | 0 – 40.6 |

*Note:* TST= total sleep time, SOL= sleep onset latency, N1= stage 1 sleep, N2= stage 2 sleep, SWS= slow wave sleep, REM= rapid eye movement sleep.

### ERP Findings

To address Hypothesis 1 (whether true memories would be associated with higher LPC amplitudes compared to false memories), the processed ERP data was analysed using a LMM predicting mean LPC amplitude from memory type and pre-stimulus activity (LPCmean ~ type*prestim + (1|ID)). Although pre-stimulus amplitude was included as a covariate in all models, only effects of variables of interest will be discussed. The LMM revealed a significant main effect of memory type on LPC amplitude (χ2(1) = 11.52, *p*<0.001, β = −2.575e^−01^), whereby amplitude was higher for true memories in contrast to false memories.

To examine Research Question 1, pertaining to the effect of sleep on the LPC amplitude for true and false memories, we predicted LPC amplitude from memory type, pre-stimulus LPC amplitude, and condition (sleep/wake) (LPCmean ~ type*prestim*condition + (1|ID)). Data were taken from delayed recognition, in order to examine the effects post sleep/wake. There was a significant interaction effect between condition and memory type on LPC amplitude (χ2(1) = 4.48, *p*<0.05, β = 0.47), whereby amplitude was higher for true memories across both conditions, and higher for both true and false memories for the nap condition in contrast to the wake condition (See Figure 3).

**Figure 3.**
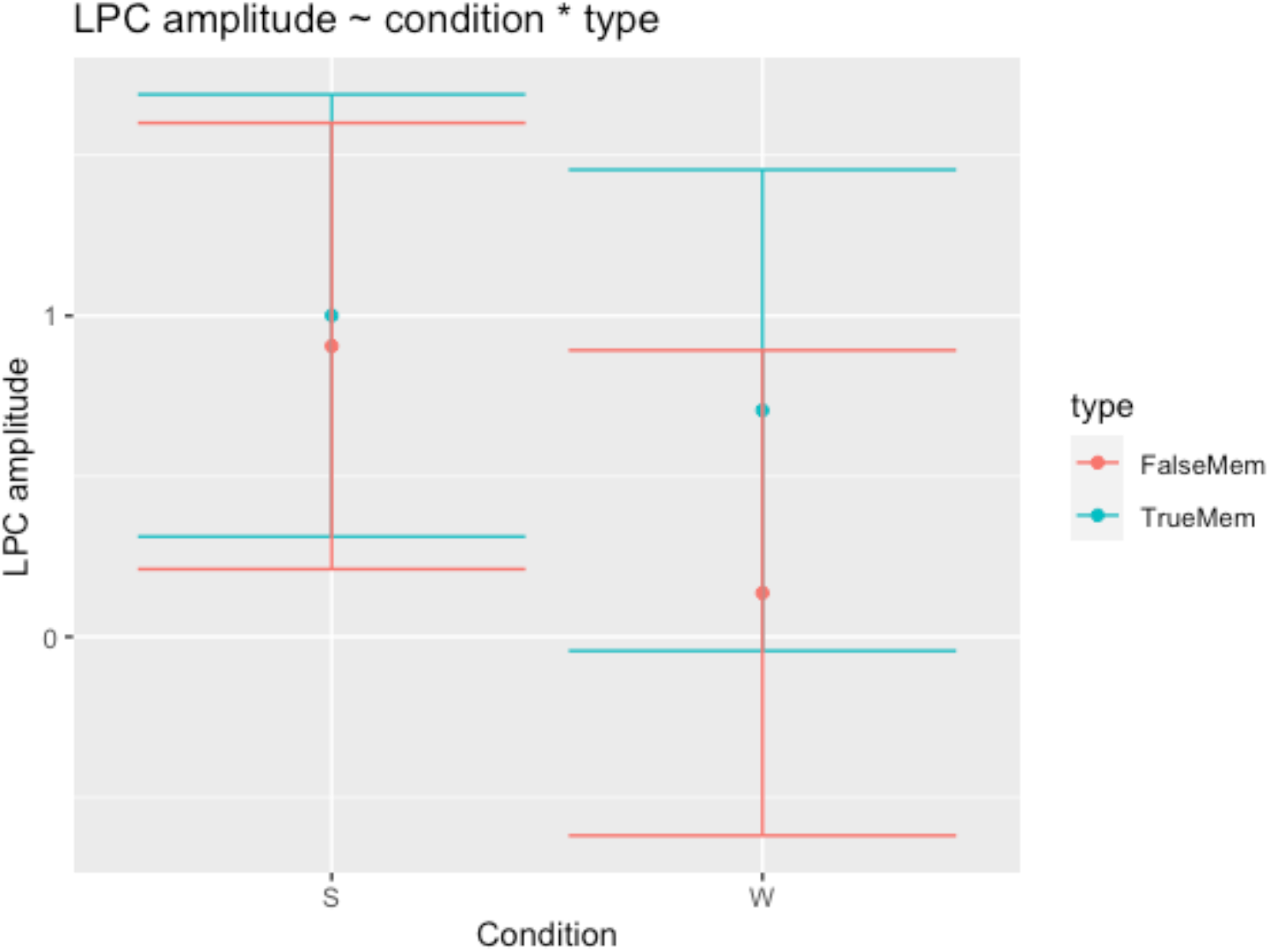
The interaction between condition and memory type on LPC amplitude, modelled over centro-medial regions. The centre dots reflect mean amplitude, and the bars indicate the lower and upper bounds of the model fit.

To examine Research Question 2 (whether P300 amplitude would differ between true and false memories), we predicted P300 amplitude from memory type, condition and pre-stimulus P300 amplitude (p300mean ~ type*prestim*condition + (1|ID)). The model detected a significant condition by memory type effect (χ2(1) = 14.52, *p*<0.001, β = −3.740e^−01^), whereby P300 amplitude was higher for false memories than for true memories across conditions, and higher for both types of memory for the nap condition, in contrast to the wake condition (See Figure 4). ERP plots for the P300 and LPC are presented in Figure 5.1, with Figure 5.2 displaying the grand average ERP over regions of interest.

**Figure 4.**
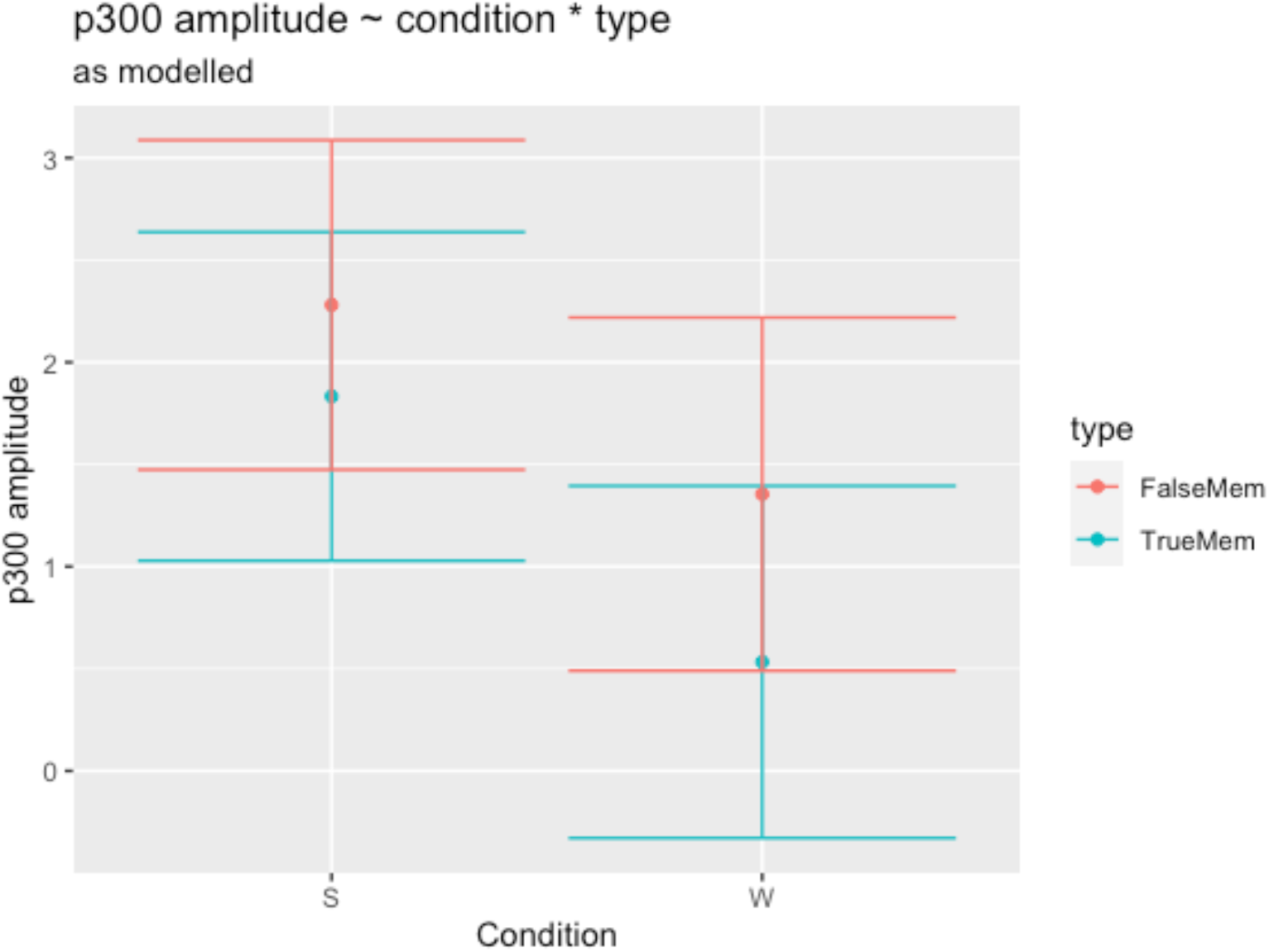
The effect of condition and memory type on P300 amplitude, modelled over posterior-medial regions. The centre dots reflect mean amplitude, and the bars indicate the lower and upper bounds of the model fit.

**Figure 5.1.**
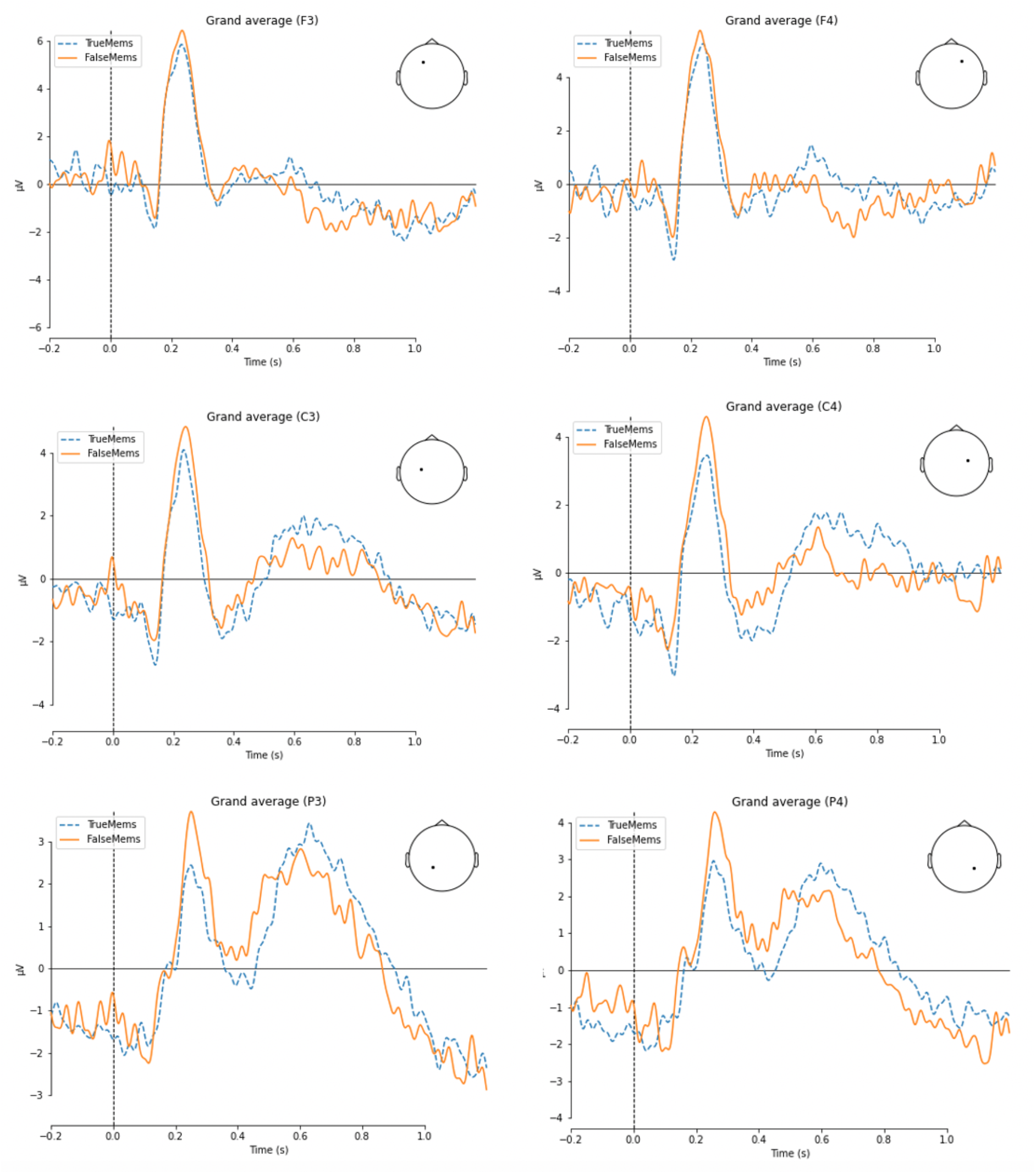
ERPs computed over electrodes F3, F4, C3, C4, P3 and P4, for both true and false memories at both immediate and delayed recognition. ERP time windows range from 200ms pre-stimulus onset, to 1 second post-stimulus onset.

**Figure 5.2.**
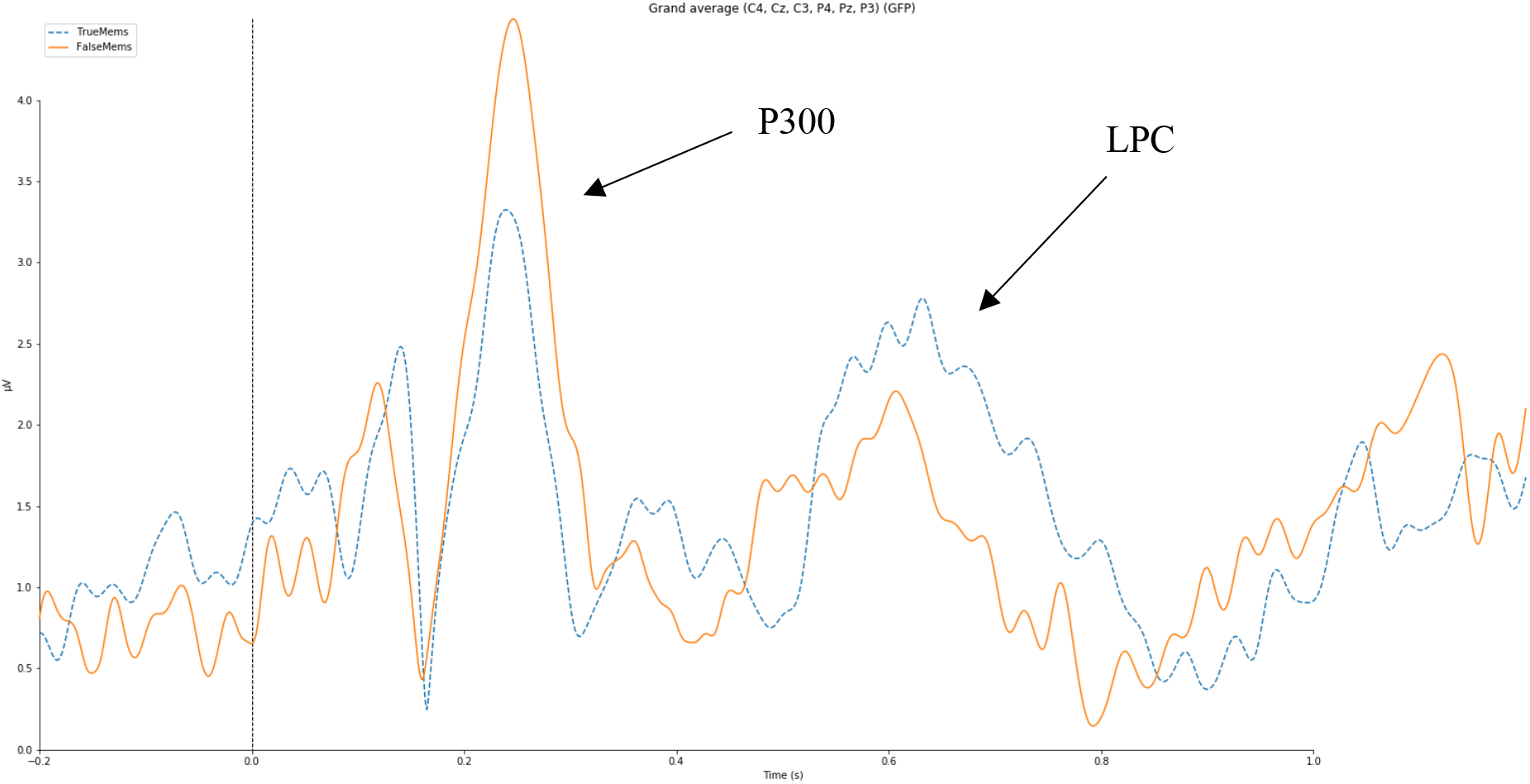
The grand average ERP, depicting a larger P300 amplitude for false, in contrast to true memories, and a larger LPC amplitude for true compared to false memories. The ERP is modelled across both immediate and delayed recognition, over regions of interest for both components (electrodes C4, Cz, C3, P4, Pz, P3).

### Combined EEG and Behavioural Modelling

In order to address Hypothesis 2 (that the nap group would have more semantic false memories than the wake group), and to examine whether LPC amplitude differences translate to behavioural effects, we predicted d’ from memory type, condition, task (IR/DR), and LPC mean amplitude (d’ ~ type*task*condition*LPCmean + (1|ID)). The effect of memory type on d’ was nonsignificant (χ2(1) = 0.00, *p*= 1.00, β = 1.419e^−03^).

To similarly examine whether P300 differences are related to behavioural differences in true and false memory endorsements, we predicted d’ from memory type, condition, task (IR/DR), and P300 mean amplitude (d’ ~ type*task*condition*p300mean + (1|ID)). The model detected a significant interaction effect of memory type, task and mean P300 amplitude on d’ (χ2(1) = 8.12, *p*<.01, β = −5.839e^−04^), whereby at immediate recognition, a larger P300 amplitude was associated with lower true memory accuracy. At delayed recognition, a larger P300 amplitude was associated with increased true memory accuracy (See Figure 6).

**Figure 6.**
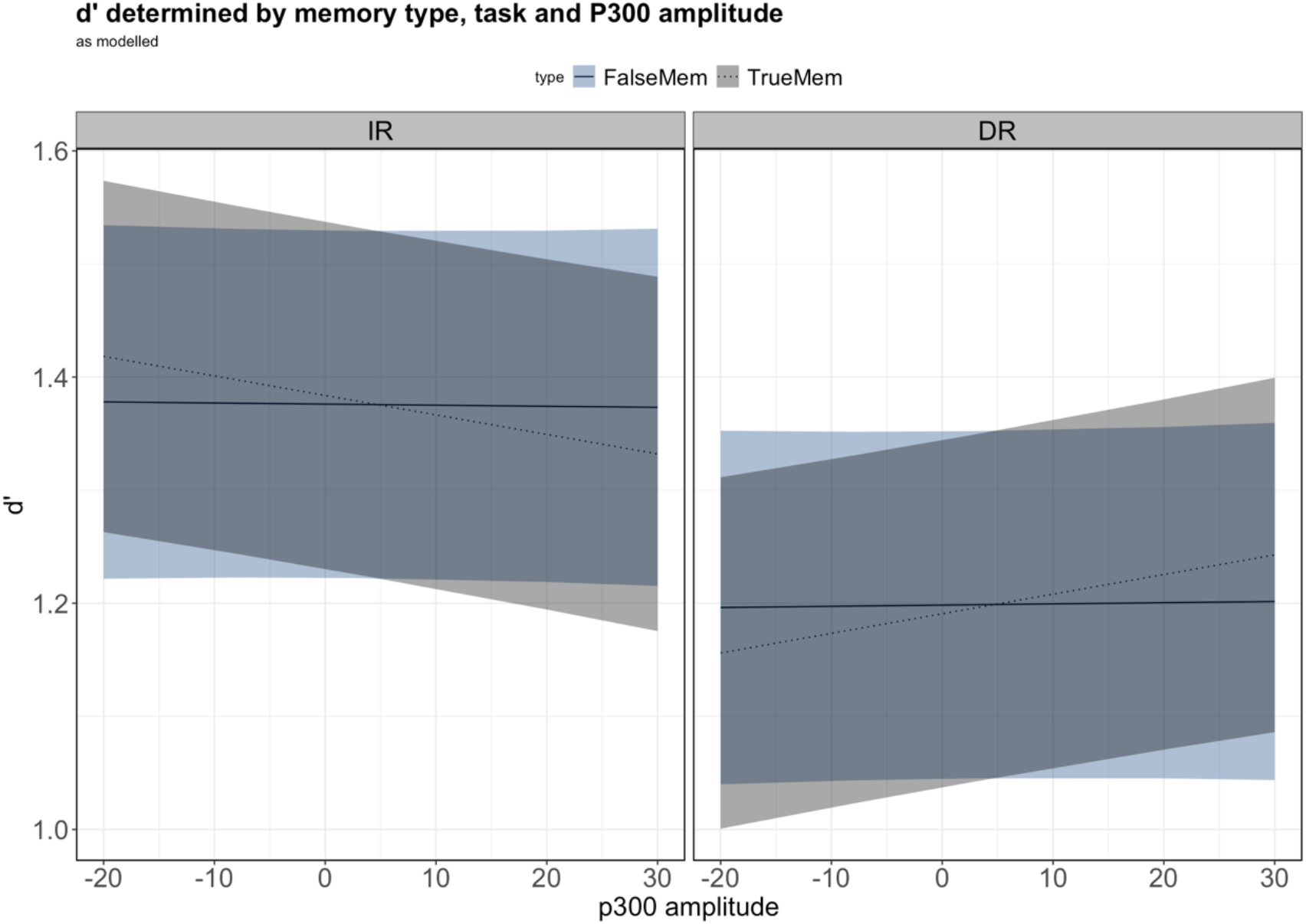
The interaction between type of memory and P300 mean amplitude on d’, faceted by task (immediate recognition; IR/delayed recognition; DR). Filled and dotted lines reflect mean d’ scores for false and true memories respectively, and shading represents 95% confidence intervals.

## Discussion

The present study explored the neurophysiological correlates of true and false semantic memories and the role of sleep in the generation thereof. As predicted, LPC amplitude was higher for true memories in contrast to false memories, independent of sleep or wake. Additionally, the interaction between memory type and condition on LPC amplitude suggests that sleep enhances true and false memory representations in the brain. Overall, P300 amplitude was higher for false than true memories, and higher for the nap group than the wake group.

The second hypothesis, that the sleep group would have more false memories than the wake group, was unsupported. However, the effect of mean P300 amplitude, memory type and task on endorsements for true and false memories suggests that neural activity and time of recognition (immediately after learning or following a delay period) produce differing outcomes for true and false semantic memory recognition.

This is the first study to demonstrate a P300 amplitude difference between semantic false memories and true memories, and to suggest that sleep has differential effects in modulating ERP components associated with true and false memories.

### The LPC as a Potential Marker for Pattern Separation

As expected, LPC amplitude was higher for true, in contrast to false memories, possibly reflecting a greater engagement of episodic recollection processes for true memories (Chen et al., 2012; Yonelinas, 2002). This finding is similar to that of previous research (Chen et al., 2012; Curran, 2000; Curran et al., 2001; Nessler et al., 2001), and provides support for the suggestion that true memories are more detailed and episodic than false memories. In extension of this finding, sleep was also found to affect this relationship, whereby the amplitude of the LPC component for both types of memory was higher following a nap than following wake, suggesting that sleep promoted the storage of true memories as distinct episodic traces.

The tendency of sleep to promote recollection-related processes could be explained by the processes of pattern separation and pattern completion (Shohamy & Turk-Browne, 2013), which suggests that the hippocampus creates distinctions between information in memory, whilst also promoting associations between related memory traces (Yassa & Stark, 2011). Pattern separation occurs when memory traces are encoded as separate representations with minimal overlap, thus facilitating retrieval of detailed episodic information with little interference (Doxey et al., 2018). The higher LPC for true memories could be indicative of pattern separation, whereby true memory-eliciting words, having been seen in the DRM learning phase, were encoded as independent, detailed memory representations, thus rendering them more episodic, and resulting in a larger LPC amplitude (Doxey et al., 2018). Additionally, as the LPC was higher for both true and false memories for the nap group than the wake group, it is possible that sleep promoted the storage of the memory traces as distinct representations, with this effect being most pronounced for true memories, likely because they have a real episodic basis.

The idea that sleep may promote pattern separation is harmonious with previous research whereby sleep was seen to stabilise pattern separation (Hanert et al., 2017). Pattern separation was also strongly correlated with hippocampal replay during sleep, prompting the suggestion that such reactivation, and potential memory consolidation, serves as an underlying mechanism for pattern separation processes (Hanert et al., 2017).

### The P300 as a Potential Marker for Pattern Completion

In addition to the suggestion that higher LPC amplitudes may be indicative of pattern completion, higher P300 amplitudes for false memories may reflect the process of pattern completion, which occurs when the hippocampus binds two items in memory together so that activation of one item prompts activation of the other (Shohamy & Turk-Browne, 2013).

As the P300 is proposed to reflect target identification and goal-directed attention (Linden, 2005), a larger P300 for semantic false memories in contrast to true memories may reflect a greater exertion of such processes, whereby the broader theme of the DRM lists became a target stimulus to which words during the recognition phase were applied and categorised accordingly. Given that semantic false memories may arise from associative processes, their presence suggests that extraction of the broader context is likely to have occurred (Straube, 2012). In extension, as critical lure words represent the broader context and are the most strongly related to it, they may have elicited a larger P300 as they were extremely salient and strongly coincided with the wider goal in mind. In contrast, study words (which elicit a true memory) vary in their relation to the broader theme, hence the smaller P300.

This process may be reflective of pattern completion, to the extent that memory traces for false memories and their broader themes were stored as overlapping representations (Yassa & Stark, 2011). Such overlap likely resulted in false memory-eliciting lures prompting memory retrieval of the overall theme of each list, making it more likely that a lure would be ‘recognised,’ because it was strongly associated with the overall theme. In extension, as the nap group exhibited larger P300 amplitudes than the wake group, it is suggested that sleep facilitated the binding of the broader list theme (the context) to the individual words (items). This may have resulted in the context/theme becoming a target, with both true and false memories being perceived as highly salient. This theory is supported by the contextual binding hypothesis, whereby SWS is proposed to drive the binding of item information and contextual information in memory (Yonelinas et al., 2019).

Although the implicit nature of such a target makes it difficult to confidently confirm that behaviourally, participants were searching for the broader theme of the lists, the existing literature provides strong evidence that implicit stimuli can act as P300-eliciting targets (Berlad & Pratt, 1995). For example, previous research has demonstrated larger P300 responses to participants’ own names, when they held another, more explicit target in mind (Berlad & Pratt, 1995; for a linguistic example, see Haupt et al., 2008).

These results may also shed light upon the discrepancies between existing ERP and false memory studies. Other ERP studies may have failed to detect a P300 difference because they lacked an appropriate retention interval between DRM learning and delayed recognition, preventing such consolidation and thematic extraction from occurring (Curran, 2000; Curran et al., 2001; Lewis et al., 2011). For example, a study by Curran and colleagues (2001) separated learning and recognition by an interval of five minutes, in contrast to the two-hour interval in the present study.

Finally, in regard to the two-process theory (Press et al, 2020), the larger P300 for false memories may be a neural reflection of a perceptual orienting towards expected stimuli. If the broader theme became a target, the larger P300 for false memory-eliciting lures may indicate a perceptual emphasis on expected stimuli, or a suppression of activity towards surprising stimuli that does not fit with the theme. This is consistent with the two-process model, which proposes that expected stimuli result in the application of a model to reduce processing effort, whilst only unexpected stimuli prompt model updating (Press et al, 2020). This is also harmonious with the traditional understanding of semantic false memories in the context of predictive coding as resulting from such model application, ultimately suggesting that the lure words must be expected, based on internal models of stimuli sets. Note that this does not necessarily imply behavioural differences as a result of schema creation, although this is an area of potential future inquiry.

In summary, ERP analyses revealed higher LPC amplitudes for true memories, higher P300 amplitudes for false memories, and higher amplitudes for both components following sleep compared to wake. This not only indicates that true and false memories have differing ERP profiles, but suggests that sleep preferentially promotes memory specificity and memory generalisation depending on whether the memory is true or false, that is, whether the memory is based on previously seen information, or on information that has an associative basis but not directly experienced. It also extends upon the idea of predictive coding, whereby related content is generalised into broader models (Aitchison & Lengyel, 2017), suggesting that such generalisation may depend on pattern separation and pattern completion processes. Predictive coding may thus be more complicated in that hippocampal neural trace replay may affect the extent to which this occurs (Yassa & Stark, 2011).

### Behavioural Findings

Contrary to our hypothesis, the nap group did not exhibit a significantly larger number of semantic false memory endorsements than the wake group. Additionally, LPC differences did not appear to translate to behavioural effects, suggesting that the episodic detail of a memory trace may not lead to better recognition.

Memory type, task, and P300 amplitude interacted to influence behavioural outcomes, such that at immediate recognition, larger P300 amplitudes were associated with fewer true memories. This pattern was reversed at delayed recognition, with higher P300 amplitudes being associated with stronger true memory accuracy. A lack of a consolidation opportunity at immediate recognition could serve as a potential explanation for this difference, whereby the short amount of time between learning and immediate recognition was not long enough to facilitate the stabilisation of the memory traces. In contrast, at delayed recognition, there had been a consolidation opportunity of the list and its broader theme, thus enabling better episodic recognition. As the P300 is associated with saliency and goal-directed target acquisition, this finding contributes to the traditional understanding of consolidation, by suggesting that the perception of target relevance in the brain changes as memories are stabilised, and that this could have implications for memory performance. The P300 may therefore serve as a potential index of consolidated vs. un-consolidated information, and future research employing a subsequent memory paradigm could verify this.

### Limitations & Future Directions

Although the present study provided novel insight into the effect of sleep on the neural representations of true and false memories, the somewhat artificial nature of the DRM renders the applicability of the present findings uncertain. Future research should investigate more complex memories to determine how well the observed effects translate to real-world situations. However, as this is one of the first studies to investigate false memories in the context of both sleep and ERPs, it is a valuable first step. Future research should also further examine the relationship between implicit target stimuli and the DRM paradigm, to obtain insight into the role of implicit stimuli in guiding memory outcomes.

Additionally, the amount of sleep the night before testing was not recorded, despite evidence that sleep before learning can influence memory (Alberca-Reina et al., 2015). Sleep microstructural variables (e.g. spindles, slow oscillations) and hippocampal neural replay were also not examined, meaning that we cannot draw strong conclusions regarding the specific sleep mechanisms that may be driving the observed effects. Future research should therefore investigate sleep microstructural variables and hippocampal replay, and record prior sleep, to differentiate between the effects of prior sleep and the effects of post-learning sleep-based consolidation.

Finally, future research may also benefit from an analysis of the semantic overlap of the DRM words, in order to better delineate the conditions that favour pattern separation or pattern completion. Previous literature suggests that the semantic overlap in content determines the likelihood of either process (Libby et al, 2018). However, the present study lacked an examination of how closely related the study words and the false memory-eliciting lures were. In future, this could contribute insight into how pattern separation and completion act to facilitate memory and learning.

## Conclusion

The present findings indicate that amplitude differences in the P300 and the LPC component differentiate between semantic false memories and true memories, and that sleep strengthens memory traces for both types of memory. The research also suggests that memory saliency may improve episodic recognition following a consolidation opportunity. This has important implications for understanding the learning process, as it suggests that learning could be enhanced according to the perceived relevance of a stimulus. Additionally, the findings further illuminate the relationship between sleep and memory overlap, with the LPC and P300 serving as potential indices of pattern separation and pattern completion, respectively. The present research may thus extend upon the traditional predictive coding perspective wherein all content is generalised into a broader model (Aitchison & Lengyel, 2017), by suggesting that memory specificity and memory generalisation within neural representations are preferentially promoted depending on the episodic basis of a memory; that is, whether the memory is true or false.

Therefore, the current study provides valuable insight into the underlying mechanisms of true and false memory formation, whilst also demonstrating the tendency of sleep to enhance neural representations for both types of memory. This ultimately provides support for the perspective of false memories as being reflective of associative, adaptive processes, rather than simply being indicative of memory errors.

The authors wish to thank Hayley Caldwell for assistance with data collection and Louise Kyriaki for editing assistance.

## Notes

### Competing Interest Statement

The authors have declared no competing interest.

## Reference List

Abe, N., Okuda, J., Suzuki, M., Sasaki, H., Matsuda, T., Mori, E., … & Fujii, T. (2008). Neural correlates of true memory, false memory, and deception. Cerebral Cortex, 18(12), 2811–2819. doi: 10.1093/cercor/bhn037.

Achermann, P., & Borbély, A. A. (1994). Simulation of daytime vigilance by the additive interaction of a homeostatic and a circadian process. Biological Cybernetics, 71(2), 115–121. Retrieved from https://link.springer.com/content/pdf/10.1007%2FBF00197314.pdf.

Aitchison, L., & Lengyel, M. (2017). With or without you: predictive coding and Bayesian inference in the brain. Current Opinion in Neurobiology, 46, 219–227. doi: 10.1016/j.conb.2017.08.010.

Alberca-Reina, E., Cantero, J. L., & Atienza, M. (2015). Impact of sleep loss before learning on cortical dynamics during memory retrieval. Neuroimage, 123, 51–62. doi: 10.1016/j.neuroimage.2015.08.033.

Allen, J. J., & Mertens, R. (2009). Limitations to the detection of deception: True and false recollections are poorly distinguished using an event-related potential procedure. Social Neuroscience, 4(6), 473–490. doi: 10.1080/17470910802109939.

Alsaadi, S. M., McAuley, J. H., Hush, J. M., Bartlett, D. J., Henschke, N., Grunstein, R. R., & Maher, C. G. (2013). Detecting insomnia in patients with low back pain: accuracy of four self-report sleep measures. BMC Musculoskeletal Disorders, 14(1), 196.

Backhaus, J., Junghanns, K., Broocks, A., Riemann, D., & Hohagen, F. (2002). Test–retest reliability and validity of the Pittsburgh Sleep Quality Index in primary insomnia. Journal of Psychosomatic Research, 53(3), 737–740. doi: 10.1016/S0022-3999(02)00330-6.

Bates, D., Mächler, M., Bolker, B., & Walker, S. (2014). Fitting linear mixed-effects models using lme4. arXiv preprint arXiv:1406.5823. Retrieved from https://arxiv.org/abs/1406.5823.

Beato, M. S., Boldini, A., & Cadavid, S. (2012). False memory and level of processing effect: An event-related potential study. Neuroreport, 23(13), 804–808. doi: 10.1097/WNR.0b013e32835734de.

Beato, M. S., & Díez, E. (2011). False recognition production indexes in Spanish for 60 DRM lists with three critical words. Behavior Research Methods, 43(2), 499–507. doi: 10.3758/s13428-010-0045-9.

Berlad, I., & Pratt, H. (1995). P300 in response to the subject’s own name. Electroencephalography and Clinical Neurophysiology/Evoked Potentials Section, 96(5), 472–474. doi: 10.1016/0168-5597(95)00116-A.

Berry, R. B., Brooks, R., Gamaldo, C. E., Harding, S. M., Marcus, C., & Vaughn, B. V. (2012). The AASM manual for the scoring of sleep and associated events. Rules, Terminology and Technical Specifications, Darien, Illinois, American Academy of Sleep Medicine, 176. Version 2.4: April 2017

Boldini, A., Beato, M. S., & Cadavid, S. (2013). Modality-match effect in false recognition: An event-related potential study. Neuroreport, 24(3), 108–113. doi: 10.1097/WNR.0b013e32835c93e3.

BrainAmp DC (38 channels) [Apparatus]. (2015). Gilching, Germany: Brain Products GmbH.

Bush, A. L., Armento, M. E., Weiss, B. J., Rhoades, H. M., Novy, D. M., Wilson, N. L., … & Stanley, M. A. (2012). The Pittsburgh Sleep Quality Index in older primary care patients with generalized anxiety disorder: psychometrics and outcomes following cognitive behavioral therapy. Psychiatry Research, 199(1), 24–30.

Buysse, D. J., Reynolds III, C. F., Monk, T. H., Berman, S. R., & Kupfer, D. J. (1989). The Pittsburgh Sleep Quality Index: a new instrument for psychiatric practice and research. Psychiatry Research, 28(2), 193–213. doi: 10.1016/0165-1781(89)90047-4.

Campanella, S., Bruyer, R., Froidbise, S., Rossignol, M., Joassin, F., Kornreich, C., … & Verbanck, P. (2010). Is two better than one? A cross-modal oddball paradigm reveals greater sensitivity of the P300 to emotional face-voice associations. Clinical Neurophysiology, 121(11), 1855–1862. doi: 10.1016/j.clinph.2010.04.004.

Carpenter, J. S., & Andrykowski, M. A. (1998). Psychometric evaluation of the Pittsburgh sleep quality index. Journal of Psychosomatic Research, 45(1), 5–13. doi: 10.1016/S0022-3999(97)00298-5.

Chatburn, A., Kohler, M. J., Payne, J. D., & Drummond, S. P. (2017). The effects of sleep restriction and sleep deprivation in producing false memories. Neurobiology of learning and memory, 137, 107–113. doi: 10.1016/j.nlm.2016.11.017.

Chen, H., Voss, J. L., & Guo, C. (2012). Event-related brain potentials that distinguish false memory for events that occurred only seconds in the past. Behavioral and Brain Functions, 8(1), 36. doi: 10.1186/1744-9081-8-36.

Curran, T. (2000). Brain potentials of recollection and familiarity. Memory & Cognition, 28(6), 923–938. doi: 10.3758/BF03209340.

Curran, T., Schacter, D. L., Johnson, M. K., & Spinks, R. (2001). Brain potentials reflect behavioral differences in true and false recognition. Journal of Cognitive Neuroscience, 13(2), 201–216. doi: 10.1162/089892901564261.

Deese, J. (1959). On the prediction of occurrence of particular verbal intrusions in immediate recall. Journal of Experimental Psychology, 58(1), 17. doi: 10.1037/h0046671.

Diekelmann, S., Born, J., & Wagner, U. (2009). Sleep enhances false memories depending on general memory performance. Behavioural Brain Research, 208(2), 425–429. doi: 10.1016/j.bbr.2009.12.021.

Diekelmann, S., Landolt, H. P., Lahl, O., Born, J., & Wagner, U. (2008). Sleep loss produces false memories. PloS one, 3(10). doi: 10.1371/journal.pone.0003512.

Doxey, C. R., Hodges, C. B., Bodily, T. A., Muncy, N. M., & Kirwan, C. B. (2018). The effects of sleep on the neural correlates of pattern separation. Hippocampus, 28(2), 108–120. doi: 10.1002/hipo.22814.

Durrant, S. J., Taylor, C., Cairney, S., & Lewis, P. A. (2011). Sleep-dependent consolidation of statistical learning. Neuropsychologia, 49(5), 1322–1331. doi: 10.1016/j.neuropsychologia.2011.02.015.

Faul, F., Erdfelder, E., Lang, A. G., & Buchner, A. (2007). G* Power 3: A flexible statistical power analysis program for the social, behavioral, and biomedical sciences. Behavior Research Methods, 39(2), 175–191. doi: 10.3758/BF03193146.

Fenn, K. M., Gallo, D. A., Margoliash, D., Roediger, H. L., & Nusbaum, H. C. (2009). Reduced false memory after sleep. Learning & Memory, 16(9), 509–513. doi: 10.1101/lm.1500808.

Fox, J. (2015). Applied regression analysis and generalized linear models. Sage Publications.

Fox, J., Weisberg, S., Adler, D., Bates, D., Baud-Bovy, G., Ellison, S., … & Heiberger, R. (2012). Package ‘car’. Vienna: R Foundation for Statistical Computing. Retrieved from https://r-forge.r-project.org/projects/car/.

Geng, H., Qi, Y., Li, Y., Fan, S., Wu, Y., & Zhu, Y. (2007). Neurophysiological correlates of memory illusion in both encoding and retrieval phases. Brain Research, 1136, 154–168. doi: 10.1016/j.brainres.2006.12.027.

Gobin, C. M., Banks, J. B., Fins, A. I., & Tartar, J. L. (2015). Poor sleep quality is associated with a negative cognitive bias and decreased sustained attention. Journal of Sleep Research, 24(5), 535–542. doi: 10.1111/jsr.12302.

Gramfort, A., Luessi, M., Larson, E., Engemann, D. A., Strohmeier, D., Brodbeck, C., … & Hämäläinen, M. (2013). MEG and EEG data analysis with MNE-Python. Frontiers in neuroscience, 7, 267. doi: 10.3389/fnins.2013.00267.

Hanert, A., Weber, F. D., Pedersen, A., Born, J., & Bartsch, T. (2017). Sleep in humans stabilizes pattern separation performance. Journal of Neuroscience, 37(50), 12238–12246. doi: 10.1523/JNEUROSCI.1189-17.2017.

Haupt, F. S., Schlesewsky, M., Roehm, D., Friederici, A. D., & Bornkessel-Schlesewsky, I. (2008). The status of subject–object reanalyses in the language comprehension architecture. Journal of Memory and Language, 59(1), 54–96. doi: 10.1016/j.jml.2008.02.003.

Heilbron, M., & Chait, M. (2017). Great expectations: is there evidence for predictive coding in auditory cortex?. Neuroscience. doi: 10.1016/j.neuroscience.2017.07.061.

Jou, J., Arredondo, M. L., Li, C., Escamilla, E. E., & Zuniga, R. (2017). The effects of increasing semantic-associate list length on the Deese–Roediger–McDermott false recognition memory: Dual false-memory process in retrieval from sub-and supraspan lists. The Quarterly Journal of Experimental Psychology, 70(10), 2076–2093. doi: 10.1080/17470218.2016.1222446.

Kompatsiari, K., Candrian, G., & Mueller, A. (2016). Test-retest reliability of ERP components: A short-term replication of a visual Go/NoGo task in ADHD subjects. Neuroscience Letters, 617, 166–172. doi: 10.1016/j.neulet.2016.02.012.

Lewis, P. A., Cairney, S., Manning, L., & Critchley, H. D. (2011). The impact of overnight consolidation upon memory for emotional and neutral encoding contexts. Neuropsychologia, 49(9), 2619–2629. doi: 10.1016/j.neuropsychologia.2011.05.009.

Lewis, P. A., & Durrant, S. J. (2011). Overlapping memory replay during sleep builds cognitive schemata. Trends in Cognitive Sciences, 15(8), 343–351. doi: 10.1016/j.tics.2011.06.004.

Libby, L. A., Reagh, Z. M., Bouffard, N. R., Ragland, J. D., & Ranganath, C. (2019). The hippocampus generalizes across memories that share item and context information. Journal of Cognitive Neuroscience, 31(1), 24–35. doi: 10.1162/jocn_a_01345.

Linden, D. E. (2005). The P300: Where in the brain is it produced and what does it tell us?. The Neuroscientist, 11(6), 563–576. doi: 10.1177/1073858405280524.

Makowski, D. (2018). The psycho package: An efficient and publishing-oriented workflow for psychological science. J. Open Source Software, 3(22), 470. doi: 10.21105/joss.00470.

Mathôt, S., Schreij, D., & Theeuwes, J. (2012). OpenSesame: An open-source, graphical experiment builder for the social sciences. Behavior Research Methods, 44(2), 314–324. doi: doi.org/10.3758/s13428-011-0168-7.

McNicol, D. (2005). A primer of signal detection theory. New York, MH. Psychology Press.

Miller, A. R., Baratta, C., Wynveen, C., & Rosenfeld, J. P. (2001). P300 latency, but not amplitude or topography, distinguishes between true and false recognition. Journal of Experimental Psychology: Learning, Memory, and Cognition, 27(2), 354–361. doi: 10.1037/0278-7393.27.2.354.

Mollayeva, T., Thurairajah, P., Burton, K., Mollayeva, S., Shapiro, C. M., & Colantonio, A. (2016). The Pittsburgh sleep quality index as a screening tool for sleep dysfunction in clinical and non-clinical samples: a systematic review and meta-analysis. Sleep Medicine Reviews, 25, 52–73. doi: 10.1016/j.smrv.2015.01.009.

Monk, T. H. (2005). The post-lunch dip in performance. Clinics in Sports Medicine, 24(2), e15–e23. doi: 10.1016/j.csm.2004.12.002.

Nessler, D., Mecklinger, A., & Penney, T. B. (2001). Event related brain potentials and illusory memories: the effects of differential encoding. Cognitive Brain Research, 10(3), 283–301. doi: 10.1016/S0926-6410(00)00049-5.

Nicholls, M. E., Thomas, N. A., Loetscher, T., & Grimshaw, G. M. (2013). The Flinders Handedness survey (FLANDERS): a brief measure of skilled hand preference. Cortex, 49(10), 2914–2926. doi: 10.1016/j.cortex.2013.02.002.

Payne, J. D., Schacter, D. L., Propper, R. E., Huang, L. W., Wamsley, E. J., Tucker, M. A., … & Stickgold, R. (2009). The role of sleep in false memory formation. Neurobiology of Learning and Memory, 92(3), 327–334. doi: 10.1016/j.nlm.2009.03.007.

Press, C., Kok, P., & Yon, D. (2020). The perceptual prediction paradox. Trends in Cognitive Sciences. doi: 10.1016/j.tics.2019.11.003.

Rasch, B., & Born, J. (2008). Reactivation and consolidation of memory during sleep. Current Directions in Psychological Science, 17(3), 188–192. doi: 10.1111/j.1467-8721.2008.00572.x.

R Core Team (2018). R: A language and environment for statistical computing. R Foundation for Statistical Computing, Vienna, Austria. Retrieved from http://www.R-project.org/.

Roediger, H. L., & McDermott, K. B. (1995). Creating false memories: Remembering words not presented in lists. Journal of Experimental Psychology: Learning, Memory, and Cognition, 21(4), 803.

Shaw, J. J., & Monaghan, P. (2017). Lateralised sleep spindles relate to false memory generation. Neuropsychologia, 107(1), 60–67. doi: 10.1016/j.neuropsychologia.2017.11.002.

Shohamy, D., & Turk-Browne, N. B. (2013). Mechanisms for widespread hippocampal involvement in cognition. Journal of Experimental Psychology: General, 142(4), 1159–1170. doi: 10.1037/a0034461.

Stadler, M. A., Roediger, H. L., & McDermott, K. B. (1999). Norms for word lists that create false memories. Memory & Cognition, 27(3), 494–500. Retrieved from https://link.springer.com/article/10.3758/BF03211543.

Straube, B. (2012). An overview of the neuro-cognitive processes involved in the encoding, consolidation, and retrieval of true and false memories. Behavioral and Brain Functions, 8(1), 35. doi: 10.1186/1744-9081-8-35.

Wickham, H. (2016). ggplot2: Elegant Graphics for Data Analysis. Springer, New York. Retrieved from https://ggplot2.tidyverse.org.

Wiese, H., & Daum, I. (2006). Frontal positivity discriminates true from false recognition. Brain Research, 1075(1), 183–192. doi: 10.1016/j.brainres.2005.12.117.

Yassa, M. A., & Stark, C. E. (2011). Pattern separation in the hippocampus. Trends in neurosciences, 34(10), 515–525. doi: 10.1016/j.tins.2011.06.006.

Yonelinas, A. P. (2002). The nature of recollection and familiarity: A review of 30 years of research. Journal of memory and language, 46(3), 441–517. doi: 10.1006/jmla.2002.2864.

Yonelinas, A. P., Ranganath, C., Ekstrom, A. D., & Wiltgen, B. J. (2019). A contextual binding theory of episodic memory: systems consolidation reconsidered. Nature Reviews Neuroscience, 20(1), 364–375. doi: 0.1038/s41583-019-0150-4.

